# Cluster Analysis of SARS-CoV-2 Gene using Deep Learning Autoencoder: Gene Profiling for Mutations and Transitions

**DOI:** 10.1101/2021.03.16.435601

**Authors:** Jun Miyake, Takaaki Sato, Shunsuke Baba, Hayao Nakamura, Hirohiko Niioka, Yoshihisa Nakazawa

## Abstract

We report on a method for analyzing the variant of coronavirus genes using autoencoder. Since coronaviruses have mutated rapidly and generated a large number of genotypes, an appropriate method for understanding the entire population is required. The method using autoencoder meets this requirement and is suitable for understanding how and when the variants emarge and disappear. For the over 30,000 SARS-CoV-2 ORF1ab gene sequences sampled globally from December 2019 to February 2021, we were able to represent a summary of their characteristics in a 3D plot and show the expansion, decline, and transformation of the virus types over time and by region. Based on ORF1ab genes, the SARS-CoV-2 viruses were classified into five major types (A, B, C, D, and E in the order of appearance): the virus type that originated in China at the end of 2019 (type A) practically disappeared in June 2020; two virus types (types B and C) have emerged in the United States and Europe since February 2020, and type B has become a global phenomenon. Type C is only prevalent in the U.S. and is suspected to be associated with high mortality, but this type also disappeared at the end of June. Type D is only found in Australia. Currently, the epidemic is dominated by types B and E.

## Introduction

The coronavirus outbreak at the end of 2019 has had unprecedented and significant consequences. Various researches have been conducted to understand the global trend of genetic alterations (Gómez-Carballa et al. 2020; Jones and Manrique 2020; Nie et al. 2020; Rochman et al. 2020). Technologies for the analysis of viruses and other genomes have been developed mainly in the field of molecular biology for basic research. High-speed gene sequencing technology has enabled the analysis of more than 40,000 cases worldwide in one year (NCBI Nucloetide Database; NCBI Virus Database). In order to understand the alternation of viral genomes while utilizing the huge amount of information, it would be helpful to conceptualize these viral mutations and visualize the spatiotemporal transition.

We have been studying the application of deep-learning autoencoder for analyzing gene sequences (Miyake et al. 2018). The feature extraction capability of autoencoder is useful for this kind of analysis. There is no need to organize the potentially characteristic sites in the gene beforehand. In our previous study of the human leukocyte antigen A (HLA-A) gene, we discovered that autoencoder can correctly represent and classify differences in HLA-A alleles (Miyake et al. 2018). Autoencoder has the potential to extract the genetic characteristics of a gene at a level close to human recognition. A brand-new method of classification could be realized.

By using a deep learning autoencoder, various analyses of genes can be performed in a limited period of time using a GPU computer, as long as the target is about tens of thousands of genes with the length of a coronavirus genome (tens of thousands of base pairs). Autoencoder does not require a gene pre-processing, such as alignment and marking of characteristic gene sequences, nor the need to prepare supervised learning data in advance. Despite this, gene types can be classified and displayed as clusters in three-dimensional space. Similar genes in sequences form a single cluster and the group can be intuitively grasped. The spatial distances between genes/clusters can serve as an indicator of genetic relationships and may contribute to a sophisticated understanding of evolutionary processes.

In this paper, we used the ORF1ab gene sequences of the new coronaviruses (collected between December 2019 and February 2021), which were obtained from the NCBI Virus and NCBI Genbank databases, to extract the self-contained features of about 30,000 genes and display them in three-dimensional space to investigate how the SARS-CoV-2 virus mutated over time.

## Methods

The Tensor-Flow library (V2.0 downgrade to V1.0) was used as Deep Learning for autoencoder. The computer was constructed in our laboratory equipped with a GPU (NVIDIA Quadoro P-6000), i7 CPU, 64GB RAM memory. OS is Windows 10 or Linax (Ubuntu) OS. The learning was usually 1 million times. The ORF1ab gene location of each gene in the NCBI nucleotide database (NCBI Genbank Database) was determined using the reference sequence (NC_045512.2).

In order to achieve high accuracy in the analysis using autoencoder, it is desirable that the length of the sample genes is uniform within a certain range. The length of the genomes in the database is difficult to use because the sequencing methods are different and not uniform. The ORF1ab gene contains the major part of non-structural proteins (15 types) and occupies more than about 2/3 of the entire viral genome. The distribution of the length of the genome or the large number of undecided sequences makes the scattered dots wider and the 3D plot harder to see. Genome samples with two or more consecutive undetermined RNA sequences were excluded from the analysis.

The nucleotide sequence data of the new coronaviruses were obtained from the NCBI Virus Database (NCBI Virus Database). The sequencing was downloaded on February 26, 2021 (the sample collection dates correspond to December 19, 2019 to February 16, 2021.). Data that were described only up to the year and data that did not specify the collection site were excluded from the analysis. Data with collection dates only up to the month were assigned a day in the middle of the month; for the two genes with no date found in December 2019, we assigned December 19. Finally, 31050 ORF1ab gene was extracted and analyzed for characteristics (cluster analysis) and time series.

Genetic analysis by autoencoder that we have already reported (Miyake et al. 2018) was used in this study. Namely, we applied the document vector method (the nucleotide sequence was replaced by a vector (4^5^ = 1,024 dimensions) with a normalized histogram of 1,024 words consisting of 5-mer tiny nucleotide sequences without alignment). In this research, the hierarchy was compressed to four layers and three dimensions. In order to visualize the obtained 3D data, we plotted them as x, y, z coordinates in 3D space. Each dot corresponds to a variant nucleotide sequence. The spatial distance from the center of the all dots plotted in 3D space was calculated and used to represent the gene profile in a time series.

Phylogenetic trees were constructed using maximum likelihood phylogenetic analysis (RAxML) with 1000 bootstraps (GENETYX ver. 15, GENETYX Co., Tokyo, Japan). Alignment of nucleotide sequences was performed using the above software.

## Results

The ORF1ab genes, extracted from the genomes of 33,915 novel coronaviruses (12/19/2019– 02/16/2021), were categorized into eight clusters in 3D space (Fig. 1). The variation of the ORF1ab gene sequence length was small, leading to the result that the separation of the clusters was clear. The 3D-compressed dots correspond to respective RNA strands as many as the number of samples used.

**Fig. 1.**
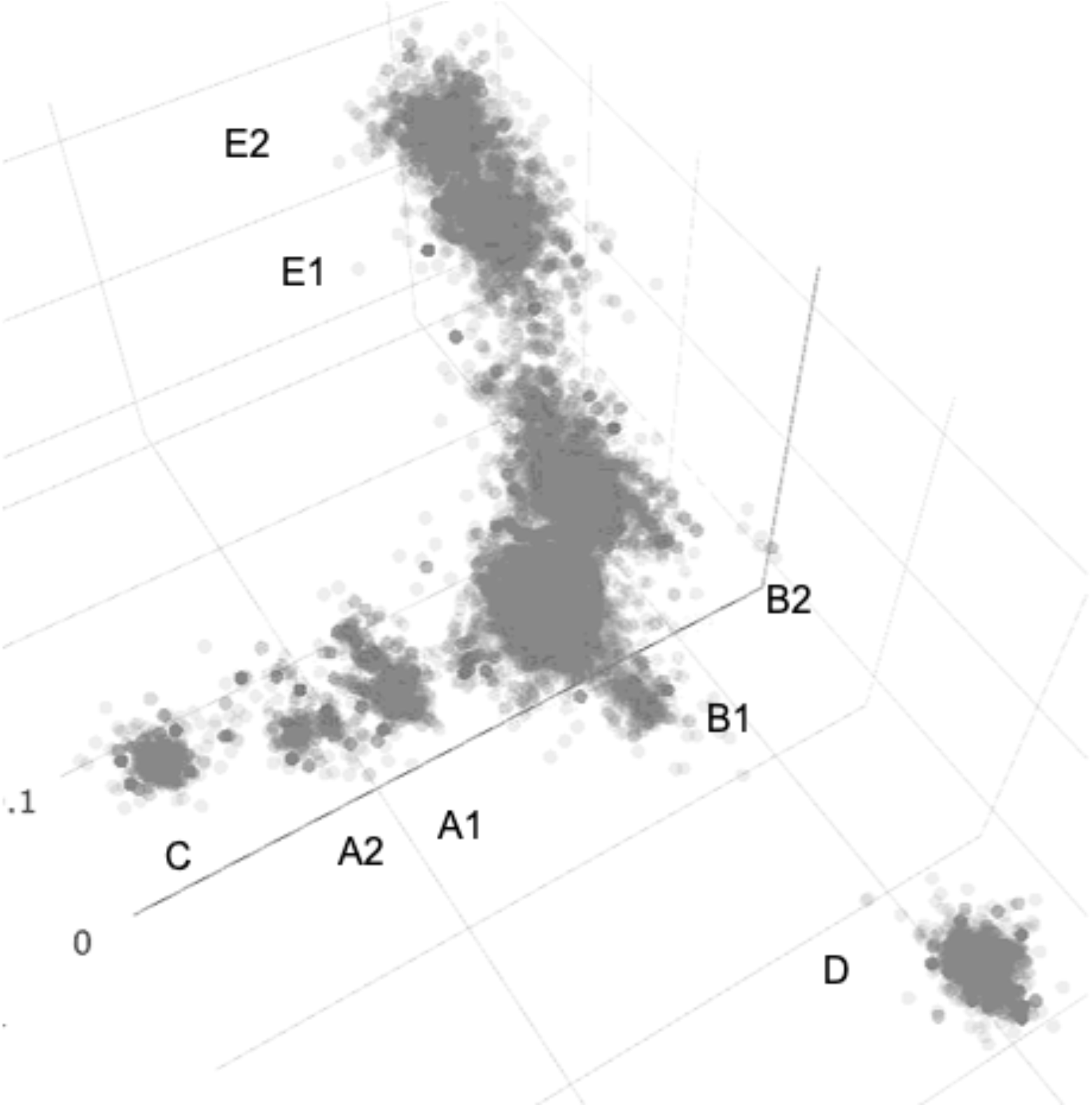
Three-dimensional plot of the ORF1ab genes. A deep-learning autoencoder classified 33,915 ORF1ab genes of SARS-CoV-2 viruses into eight clusters. The symbols of the clusters are given in the order of the time of emergence. Two clusters that appeared at the same time differed with suffix. ORF1ab genes were dissected from 33,915 genome sequences of SARS-CoV-2 viruses collected from December 2019 to February 2021. Occurrence time was shown by colored dots monthly (December 2019–February 2020) or bimonthly (March 2020–February 2021).

These dots are plotted at different spatial locations, but instead of being simply distributed, several types of sets (clusters) appear. The ORF1ab genes were shown to form the eight clusters with similar characteristics in 3D coordinates and distances from the center. Close proximity of three pairs of neighboring clusters suggested their similarities in mutation profiles, respectively. The eight clusters of the ORF1ab genes were categorized into five major groups (Fig. 1). These clusters were named A, B, C, D, and E in the order of appearance.

In order to investigate the temporal changes, we replotted the 3D dots monthly or bimonthly for the collection period (Fig. 2). The ORF1ab genes collected during the two months of December 2019 and January 2020 showed a predominance of type A cluster in the center of 3D plotted genes (Fig. 2 **a, b**). Type B became the dominant genotype from February to March, and type E became the dominant genotype from April to May. Type C started in February and fell and disappeared in June. Type D appeared in June-July and disappeared in October.

**Fig. 2.**
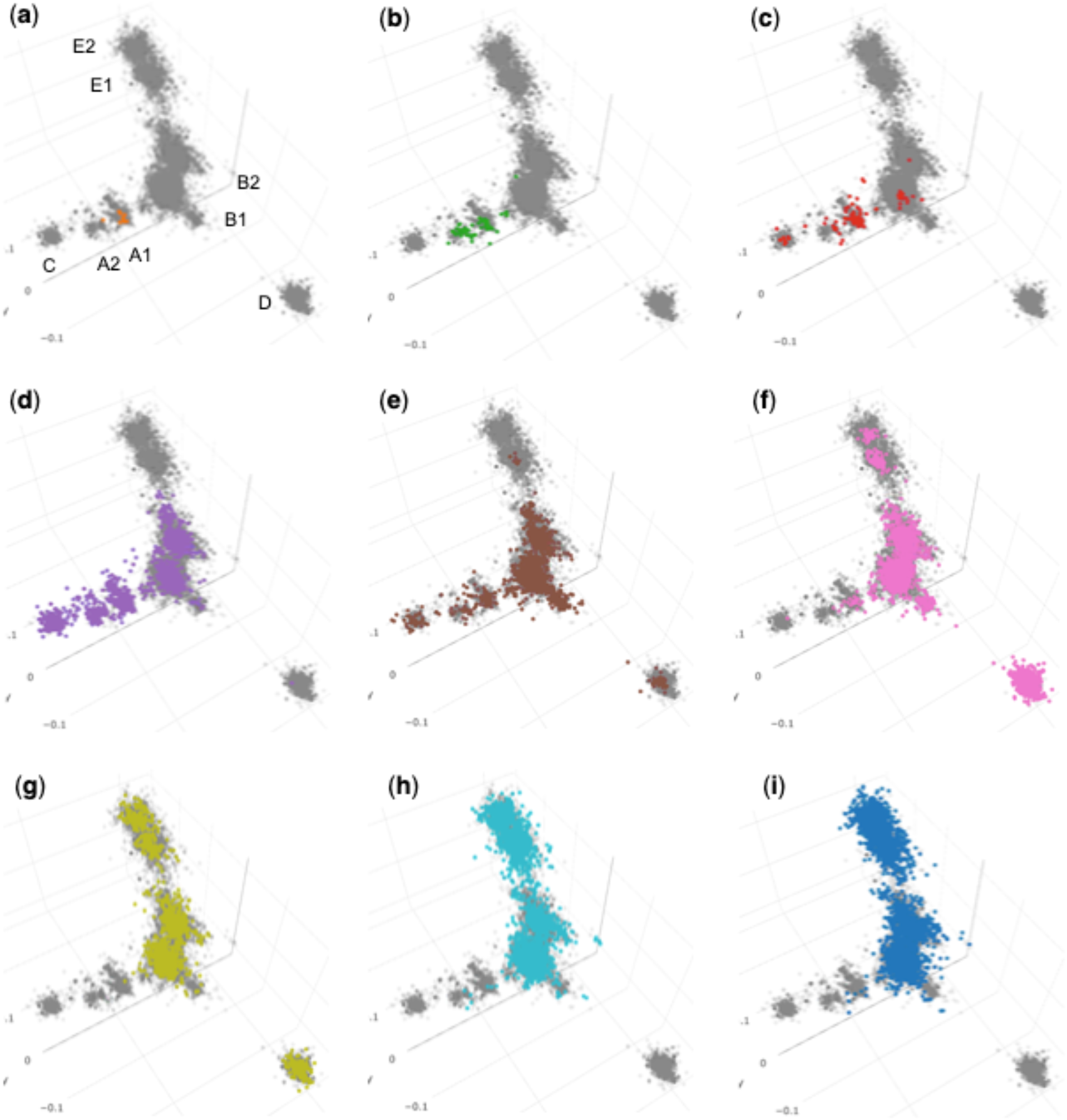
Monthly or bimonthly trend of gene clusters. (**a**) ORF1ab genes of SARS-CoV-2 viruses collected in December 2019 (orange). (**b**) January 2020 (green). (**c**) February 2020 (red). (**d**) March–April 2020 (purple). (**e**) May–June 2020 (brown). (f) July–August 2020 (pink). (**g**) September–October 2020 (yellow green). (**h**) November–December 2020 (cyan). (**i**) January–February 2021 (blue). Shown as background in light gray is the ORF1ab gene for the entire period.

The time series of the type C obtained by autoencoder analysis seems to be consistent with the emergence and disappearance and geo location of coevolving variant group 4 (CEVg4) reported by Chan et al. (Chan et al. 2020). Based on genome frequencies and geo locations, our classification of types A1, A2, B1, and D seemed to correspond to the wild type, CEVg3, CEVg1, and CEVg6, respectively. The B2 cluster is in a different location from the B1 cluster and, is a group of similar size to the B1 cluster (Figs. 1 and 2). In contrast, there is no CEVg similar to CEVg1.

The distance of each dot from the center of the all dots in the 3D space was calculated and used to represent the genotype profile in a time series by country/region. The data were color-coded by cluster and displayed separately by geographic region (Figs. 3 and 4).

**Fig. 3.**
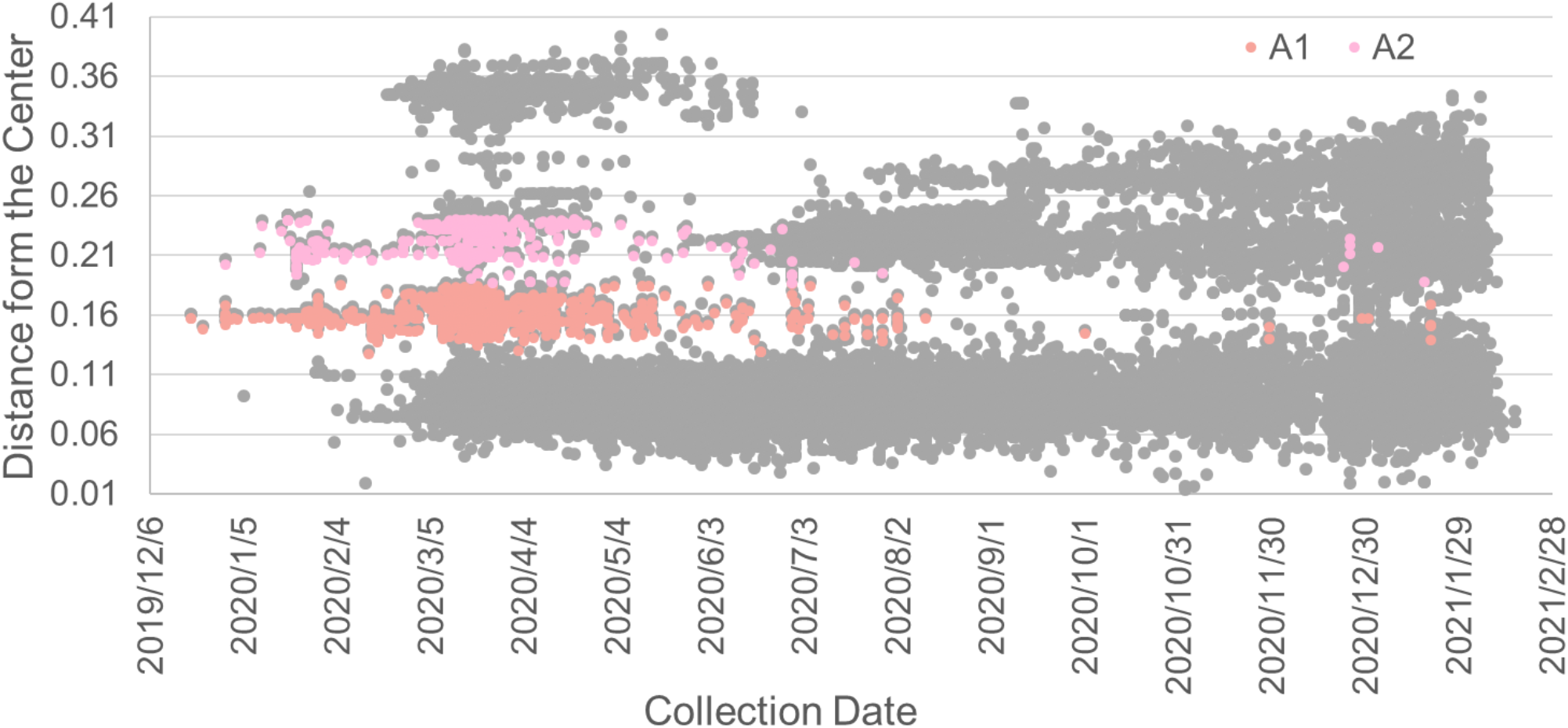
Time course plot of the SARS-CoV-2 ORF1ab gene for A1 and A2 clusters. This figure shows the way of visualization of temporal transitions in target gene clusters. The gray or colored dots, which represent the ORF1ab genes classified by autoencoder, were plotted by the collection date and distance from the center in the 3D plot in Fig. 1. Based on the coordinate data of each gene, the time evolution of a particular cluster can be shown by coloring. Here we show an example of A1 and A2 extracted.

**Fig. 4.**
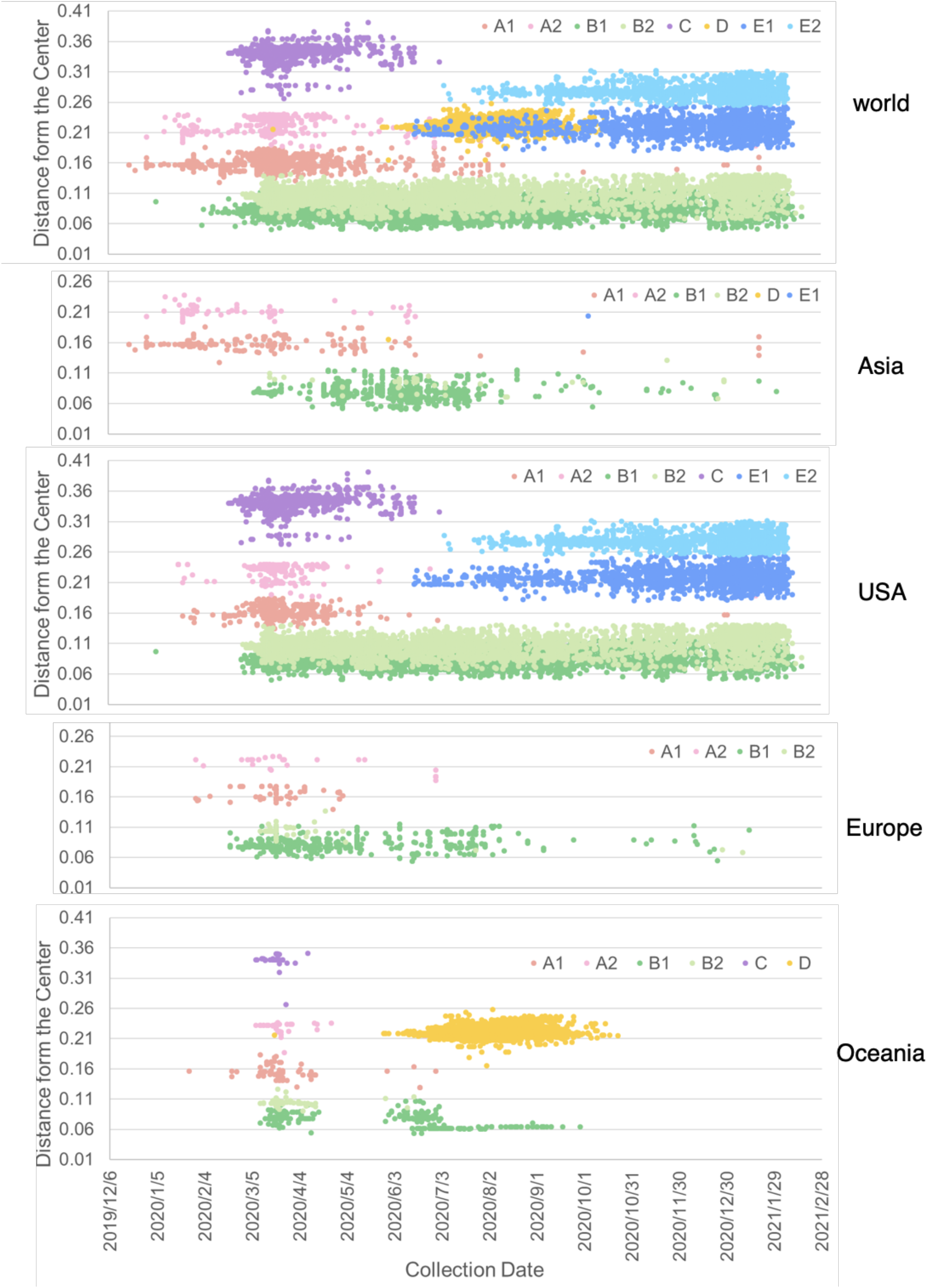
Emergence and transition of clusters. Autoencoder-classified ORF1ab genes for World, Asia, USA, Europe and Oceania are separately illustrated. Clusters are extracted as shown an example in Fig. 2. Dot colors represent different clusters (A1, red; A2, pink; B1, green; B2, light green; C, purple; D, orange; E1, blue; E2, light blue). To prevent mixing, dots in border regions between the clusters were omitted. Along the axis of the distance from the center, B2 cluster had some overlap with B1 cluster.

The stretching and extinction of genotypes was quite frequent, with a new species emerging and disappearing approximately every two months. It is unclear whether this was derived from a single species, or whether a species that originally existed was grown.

The maximum likelihood phylogenetic trees of 88 ORF1ab genes and their corresponding full-length genomes are shown in Fig. 5a and b, respectively. Genes corresponding to genotypes A1, A2, B1, B2, C, D, E1, and E2 in both the ORF1ab and full-length genomes of the phylogenetic tree are represented by the same colors as in Fig. 4. Genes classified into clusters can be regarded as having a certain degree of correlation. This is a fairly good correlation considering the fact that they have different principles.

**Fig. 5.**
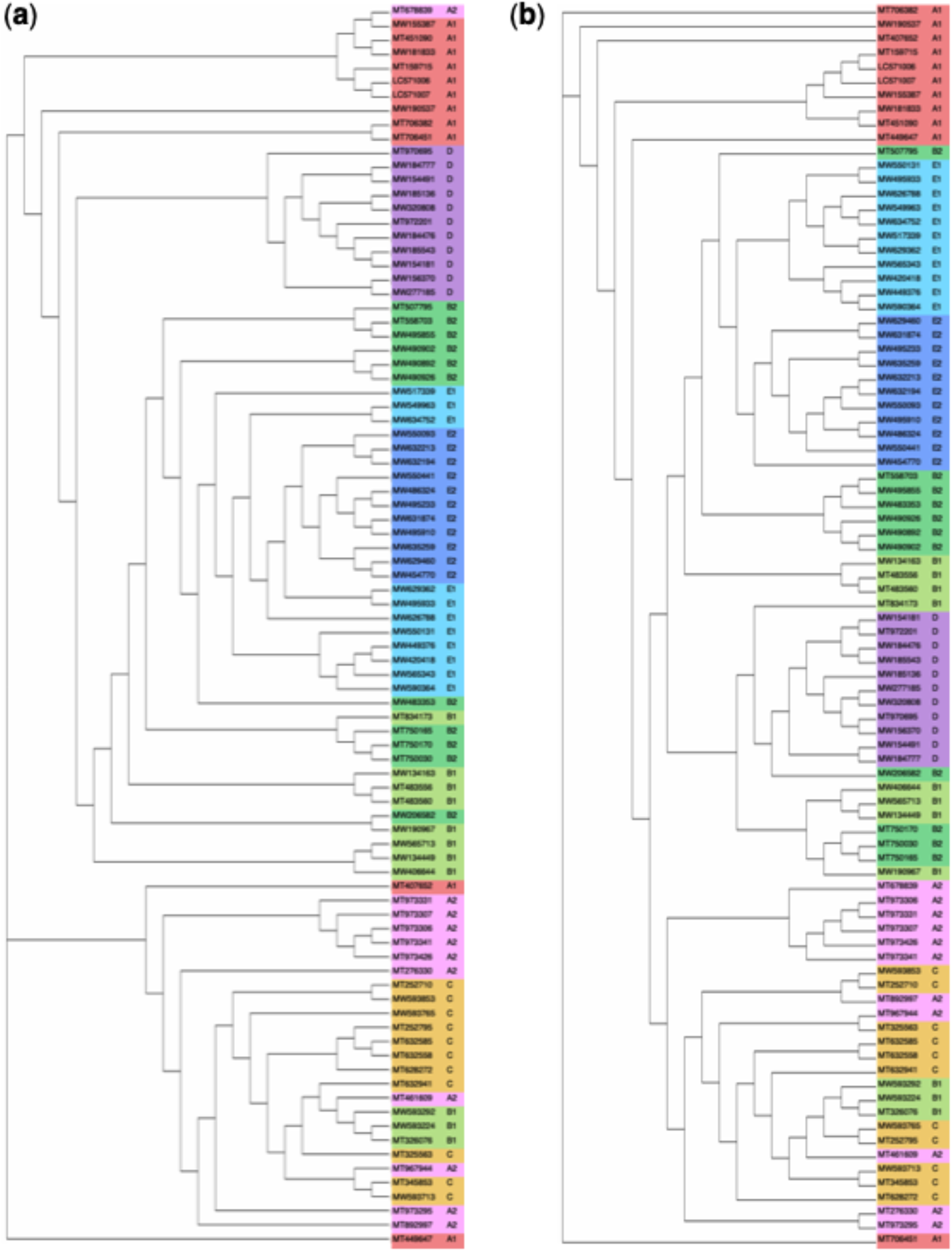
Phylogenetic trees of SARS-CoV-2 ORF1ab genes and whole genomes. (**a**) Phylogeny of 88 ORF1ab genes of SARS-CoV-2 viruses. (**b**) Phylogeny of 88 whole genomes of the SARS-CoV-2 viruses. Eleven genes from each of eight clusters were randomly selected from the central region of the respective clusters of A1, A2, B1, B2, C, D, E1 and E2 in the autoencoder 3D plot (Fig. 1). The accession numbers of the viruses used were identical between the ORF1ab genes and whole genomes. Phylogenies showed similar features between ORF1ab genes and whole genomes that members from single or two clusters formed isolated or mixed clades. Phylogenetic trees were constructed with maximum likelihood method with RAxML, 1,000 bootstraps (GENETYX ver. 15).

## Discussion

The deep-learning autoencoder was able to successfully classify the genotypes of SARS-CoV-2 viruses, and the autoencoder method is useful for overarching classification, which is similar to human cognitive abilities. It is widely recognized that the genes of coronaviruses change one after another, and the autoencoder method is a useful method for easily recognizing time-series changes. As shown in Figs. 2–**4**, it is a simple and straightforward method that allows us to grasp the elongation and disappearance of clusters in the viral genome.

Judging from the present analysis, the occurrence and disappearance of new species appears to be observed about every two months. Such correlations are available for understanding: type A first appeared in December, 2019, but largely disappeared by the end of June, 2020. The most widespread strains globally appear to be types B and E. Type B also appears to have started close to type A and spread, possibly as a result of successive changes in each.

Coronaviruses are RNA viruses and are particularly rapidly-mutated genomes. As shown in the eight clades in Fig. 1, mutations do not cause simple spread, but lead to the formation of clusters. Mutant species that pop out of the clusters form new clusters there as well. We can read a form of repeated expression and flourishing of new species in nature.

The cluster classification by the autoencoder method (shown in Fig. 1) showed a certain correlation with the classification by the phylogenetic tree method (Fig. 5). In both ORF1ab and the whole genome, a certain degree of cohesion was observed for genotypes A1, A2, B1, B2, C, D, E1, and E2, and we judged that there is considerable correlation in gene sequencing. Because both classification by autoencoder and phylogenetic tree analysis based on sequence homology and differences, the methods are do not always match perfectly in principle, but they help each other to understand classification. As shown in Fig. 5, they can be considered as essentially distant correlations as classification methods for gene sequences.

We found the eight clusters using over 30,000 SARS-CoV-2 ORF1ab genes in the NCBI Virus database, whereas Chan et al. identified nine CEVg using 86,450 genomes in the GISAID database (Chan et al. 2020). Yet there is not enough data to rigorously compare the differences between the ABCDE and CEVg classifications. autoencoder-based classification is considered to be a useful method for scanning the entire SARS-CoV-2 virus for variations or for rapid genetic classification of viral genes and viruses with a certain genetic distance.

With regard to the new coronavirus, more than 40,000 gene sequencings were performed in one year for the whole world. In order to take advantage of the vast information space made possible by next-generation sequencing technology, we believe that we need technology to grasp the entire picture of genetic variation and its distribution patterns. We hope that artificial intelligence will contribute to the development of methods for rapid recognition and classification of genetic mutations. We believe that being able to explain the direction of mutations and the principles that constrain them will make a significant contribution to this field. A better understanding of viral evolution will allow us to respond more effectively and quickly to pandemics.

We are currently working on a detailed analysis of the internal structure of the autoencoder cluster and would like to point out that there may be new applications for classification.

## Author contributions

JM designed the project; JM and TS wrote the manuscript; SB developed computer system and software; HM collected data; HN supervised the study of artificial intelligence; YN realized the project and scientifically supervised the work.

## Acknowledgements

We should like to express our thanks to Yuta Nitada, Keita Fukuda and Takuto Shimazaki of Osaka University for programming and computer operations. This work was supported partially by Global Center for Medical Engineering and Informatics of Osaka University, Hitz Research Alliance Laborator of Osaka University, and Japan Agency for Medical Research and Development (Grant Number 20bm0804008h0004.PI. Prof. S. Miyagawa of Medical School of Osaka University).

